# Transcription arrest induces formation of protective RNA granules in mitochondria

**DOI:** 10.1101/2024.09.25.614902

**Authors:** Katja G. Hansen, Autum Baxter-Koenigs, Caroline A. M. Weiss, Erik McShane, L. Stirling Churchman

**Author notes:** Corresponding author: L. Stirling Churchman, Department of Genetics, Blavatnik Institute, Harvard Medical School, Boston, MA 02115, USA.

## Abstract

Mitochondrial gene expression regulation is required for the biogenesis of oxidative phosphorylation (OXPHOS) complexes, yet the spatial organization of mitochondrial RNAs (mt-RNAs) remains unknown. Here, we investigated the spatial distribution of mt-RNAs during various cellular stresses using single-molecule RNA-FISH. We discovered that transcription inhibition leads to the formation of distinct RNA granules within mitochondria, which we term inhibition granules. These structures differ from canonical mitochondrial RNA granules (MRGs) and form in response to multiple transcription arrest conditions, including ethidium bromide treatment, specific inhibition of the mitochondrial RNA polymerase, and depletion of the SUV3 helicase. Inhibition granules appear to serve a protective function, stabilizing certain mt-mRNAs during prolonged transcription inhibition. This phenomenon coincides with an imbalance in OXPHOS complex expression, where mitochondrial-encoded transcripts decrease while nuclear-encoded subunits remain stable. We found that cells recover from transcription inhibition via resolving the granules, restarting transcription and repopulating the mitochondrial network with mt-mRNAs within hours. We suggest that inhibition granules may act as a reservoir to help overcome OXPHOS imbalance during recovery from transcription arrest.

## Introduction

Mitochondria fulfill numerous essential functions to maintain cellular homeostasis. One of the key functions is ATP production through oxidative phosphorylation (OXPHOS) via the OXPHOS complexes. Dysfunction of OXPHOS is associated with diseases, such as encephalopathies, Leigh Syndrome, cancer (Thompson Legault *et al*., 2015; Reznik *et al*., 2017; Ghezzi and Zeviani, 2018) and plays a key role in cellular aging (Houtkooper *et al*., 2013; Lima *et al*., 2022).

Mitochondria contain their own genome which is present in between 100 to 1000s copies per cell. The mitochondrial DNA (mtDNA) encodes for 13 OXPHOS subunits, 22 tRNAs, and 2 rRNAs; the residual mitochondrial proteome, including the remaining OXPHOS subunits, is nuclear-encoded and needs to be imported. In order to build functional OXPHOS complexes, gene expression has to be coordinated and balanced across compartments(Gustafsson, Falkenberg and Larsson, 2016; Isaac, McShane and Churchman, 2018; Soto *et al*., 2022).

Mitochondrial gene expression is regulated by numerous factors, allowing for the differential control of individual mtDNA molecules within the mitochondrial network. Despite the high copy number of mtDNA in cells, most mitochondrial nucleoids are tightly compacted (Brüser, Keller-Findeisen and Jakobs, 2021; Isaac *et al*., 2024), inactive for transcription (Brüser, Keller-Findeisen and Jakobs, 2021) or replication (Lewis, Uchiyama and Nunnari, 2016). The transcriptional state varies across the mitochondrial network (Brüser, Keller-Findeisen and Jakobs, 2021). Additionally, the mitochondrial RNA polymerase POLRMT can be reversibly inhibited by binding to the non-coding 7S RNA. This mechanism is thought to enable fine-tuned temporal and spatial control of POLRMT activity across different mtDNA molecules in the network (Zhu *et al*., 2022). Together, these regulatory mechanisms allow for dynamic and localized control of mitochondrial gene expression.

Similar to the nucleolus, mitochondrial RNA granules (MRGs) serve as specialized sites for RNA processing and ribosome assembly within mitochondria (Antonicka and Shoubridge, 2015; Jourdain *et al*., 2015; Popow *et al*., 2015; Tu and Barrientos, 2015; Boehm *et al*., 2017; Rey *et al*., 2020; Ohkubo *et al*., 2021; Xavier and Martinou, 2021). MRGs contain unprocessed polycistronic RNAs and key RNA binding proteins (RBPs) such as GRSF1 and FASTKD2 Within MRGs, nascent RNA is processed into mRNAs, tRNAs and rRNAs (Antonicka *et al*., 2013; Jourdain *et al*., 2013, 2015, 2016; Antonicka and Shoubridge, 2015; Popow *et al*., 2015; Rey *et al*., 2020). In addition to MRGs, other distinct RNA-containing structures have been identified within mitochondria. These include D-foci, which are sites of RNA degradation and contain the mitochondrial exonuclease, a dimer formed by the exoribonuclease PNPT1 (PNPase) and the DNA helicase SUV3 (Borowski *et al*., 2013; Xavier and Martinou, 2021), and granules containing double stranded RNA (dsRNA) (Dhir *et al*., 2018). Mitochondria may also form stress-induced RNA granules, similar to the well-characterized cytoplasmic stress granules that sequester translation initiation complexes and mRNAs during cellular stress (Anderson and Kedersha, 2009). Recent studies show the formation of mitochondrial stress granules or stress granule-like structures during certain conditions (Sun, Van Gilst and Crowder, 2023; Begeman *et al*., 2024). However, the full range of stress conditions that induce these mitochondrial structures, their RNA composition, and their functions remain to be fully elucidated.

In this study, we investigated the formation of stress-induced RNA granules inside mitochondria. Using single-molecule RNA fluorescence in situ hybridization (RNA-FISH), we examined the spatial distribution of mt-RNAs under various stress conditions. Our findings reveal that transcription inhibition triggers the formation of novel RNA granules, which we have termed “inhibition granules.” These structures are distinct from the canonical mitochondrial RNA processing granules (MRGs). We demonstrate that all mt-mRNAs localize to these inhibition granules, and importantly, these structures appear to stabilize a subset of the transcripts. Despite this protective effect, overall mt-RNA abundance decreases during transcription inhibition, leading to an imbalance in the expression of OXPHOS components. Interestingly, cells can recover from transcription inhibition by reinitiating transcription and redistributing mt-mRNAs throughout the mitochondrial network within hours upon stress release. Based on these observations, we hypothesize that inhibition granules serve as temporary RNA reservoirs which may help to restore OXPHOS balance and mitochondrial function upon stress alleviation.

## Results and discussion

### Ethidium bromide treatment leads to the formation of RNA granules

In the cytosol, stress granules (SG) form in response to stress, a process that is poorly understood in mitochondria. Therefore, we aimed to investigate the spatial distribution of mt-mRNAs during steady-state and stress conditions by using single molecule RNA-FISH. However, some mt-mRNAs are only a few hundred base pairs long (e.g. *MT-ND3* is 346 nucleotides long), rendering their visualization difficult. Hence, we employed SABER-FISH (signal amplification by nucleotide exchange reaction) (Kishi *et al*., 2019) to detect mt-mRNAs. SABER-FISH utilizes nucleotide exchange reactions to elongate primer sequences in an oligo pool creating adapters of several hundred base pairs in length. This drastically increases the detectable signals as a high number of small fluorescent oligos are associated with the adaptors (Kishi *et al*., 2019). We robustly detect mitochondrial-encoded RNAs in U2-OS cells using SABER-FISH (**Fig. 1A,B**, **EV 1A**). Our data shows that mt-mRNAs are highly abundant and distributed throughout the mitochondrial network.

**Figure 1:**
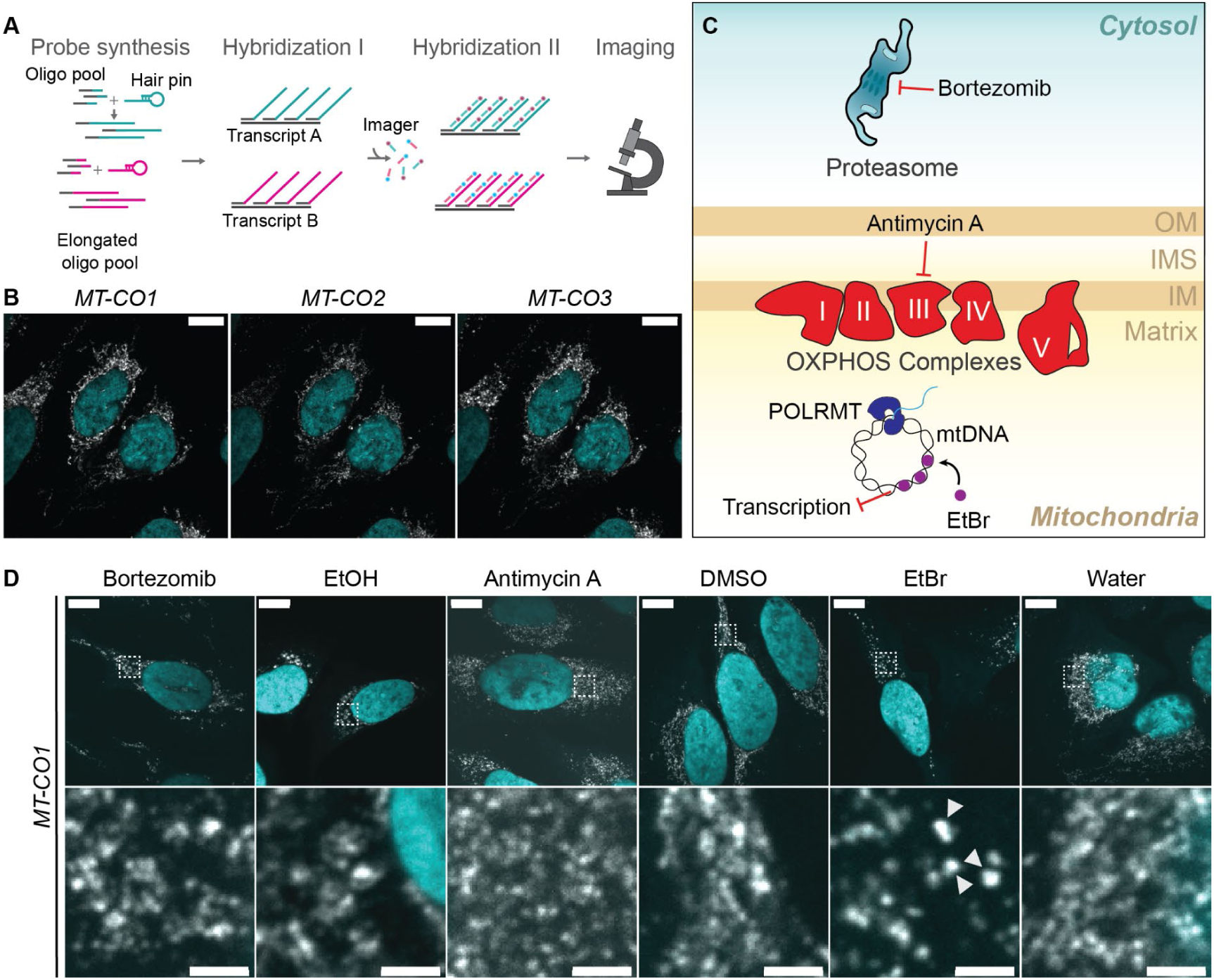
Ethidium bromide treatment leads to changes in the spatial distribution of the mitochondrial transcript *MT-CO1*. **(A)** Work-flow of SABER-FISH. Oligo pools are elongated with the help of hairpins creating long overhangs. Pink and turquoise color represent different sequences. In a first hybridization probes bind to transcripts of interests. In a second hybridization, imager probes with fluorophores bind to the complementary overhangs followed by imaging. **(B)** Confocal fluorescence microscopy of multiplexed SABER-FISH of three different transcripts in U2-OS cells. The complex IV subunit transcripts *MT-CO1*, *MT-CO2* and *MT-CO3* were labeled. The scale bars represent 10 μm. **(C)** Scheme showing acting sites of the used drugs. Bortezomib inhibits the proteasome, antimycin A blocks OXPHOS complex III and ethidium bromide (EtBr) intercalates into the mitochondrial DNA and subsequently inhibits transcription. **(D)** SABER-FISH for *MT-CO1* after different drug treatments. Cells were treated with either 100 μM Bortezomib, antimycin A or 2 μg/ml EtBr for 5 h. The respective vehicle control was used and is shown next to the corresponding drug. Images of representative cells are shown. White boxes indicate areas that are shown enlarged underneath. Scale bars of full cell images represent 10 μm and of zoom-in 2 μm. White arrows mark clustered RNA in EtBr treated cells. For all images shown in this study if not otherwise mentioned, single z-planes were chosen. DAPI staining is shown in cyan and RNA staining in gray. All images in this figure were adjusted to represent the spatial distribution and not differences in intensities.

Upon successful implementation of the SABER-FISH technology on all mt-mRNAs, we aimed to investigate possible changes in their spatial distribution during cellular stress. To cover a broad range of various stress inducers, we analyzed mt-mRNA localization after treatment with: the proteasomal inhibitor Bortezomib known to cause stress granule formation in the cytosol(Fournier, Gareau and Mazroui, 2010), the complex III inhibitor antimycin A, ethidium bromide, which intercalates into mtDNA and inhibits DNA replication and transcription and chloramphenicol, a mitochondrial translation inhibitor (**Fig. 1C**, **EV1B-D**). We treated U2-OS cells for 5 hours (**Fig. 1C**, **EV1 C, D**) or 24 hours (**Fig. EV1B**) and probed for *MT-CO1*. Of these treatments, only EtBr led to a profound visible change in RNA localization compared to its control, where *MT-CO1* formed clusters resembling RNA granules (**Fig. 1D**). The formation of RNA clusters was robust across various concentrations of EtBr (**Fig. EV1D**) and occurred also in HeLa cells (**Fig. EV 1E**).

### RNA granules form under multiple mitochondrial transcriptional perturbations

To further characterize the EtBr-induced granules, we conducted a comprehensive analysis of all mitochondrial-encoded OXPHOS subunits. Using RNA-FISH probes specific to each mitochondrial transcript, we observed that all mitochondrial mRNAs exhibited a granular distribution pattern similar to that of *MT-CO1*. This consistent behavior across all mitochondrial transcripts suggests a global response of mitochondrial RNAs to transcription inhibition, rather than a transcript-specific phenomenon (**Fig. 2A**, **EV 1C**). To investigate the timing of granule formation, we performed a time course experiment, which showed that the first granules were visible after 2 hours of EtBr treatment (**Fig. 2B**). To quantify RNA granule formation after EtBr treatment, we measured the fraction of cellular area occupied by RNA, calculated as the segmented area of the RNA signal as a fraction of the total cell area, a measure used in the analysis of granule formation in the cytosol(Mateju *et al*., 2020) (**Fig. 2C**). The fraction of cellular area occupied in the EtBr treated samples decreased for the majority of the transcripts, indicating that RNAs are less distributed and form more granules, in both biological replicates (**Fig. 2D**, **EV 2A**).

**Figure 2:**
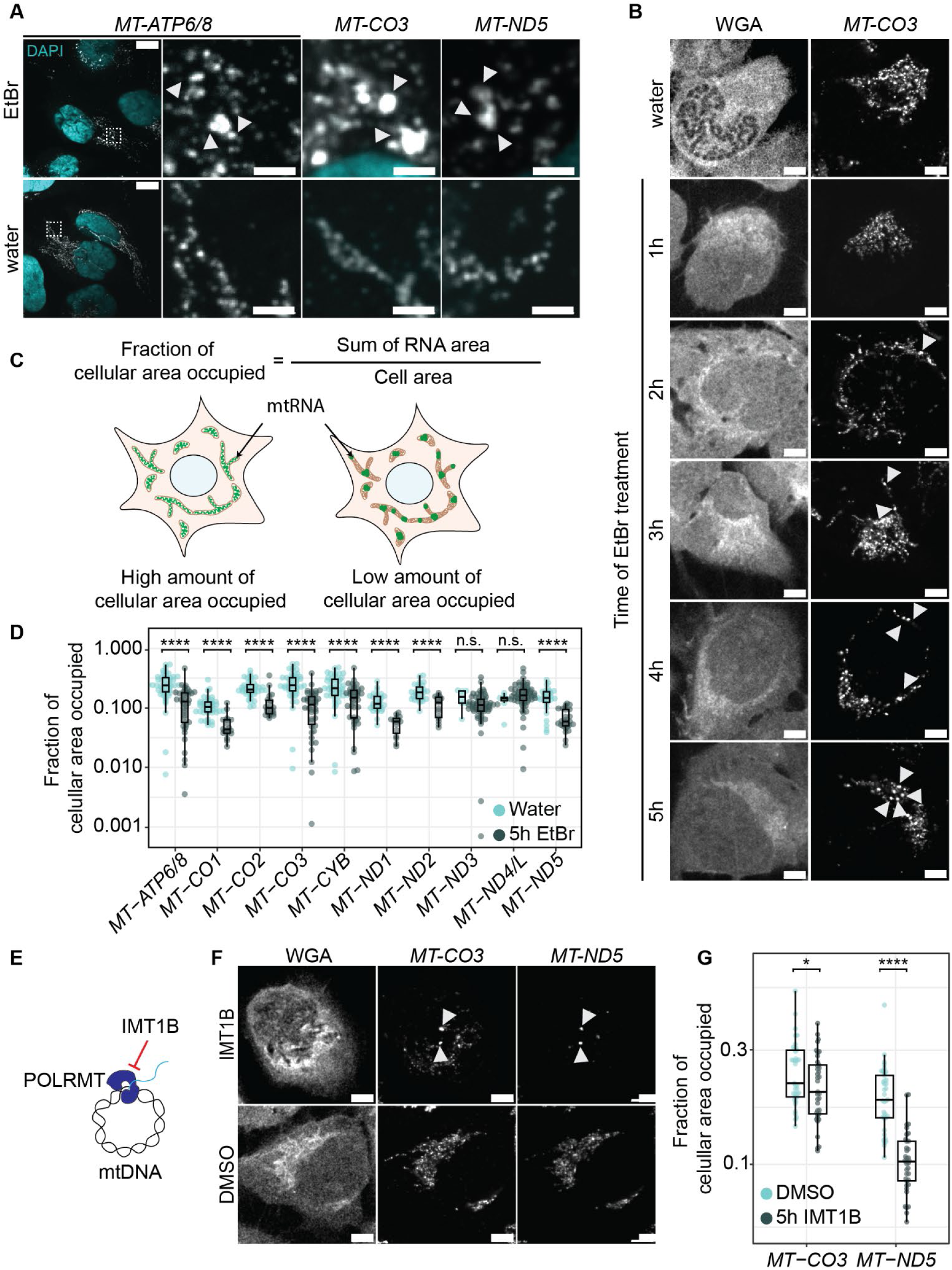
Transcription arrest leads to the formation of RNA granules. **(A)** Example transcripts forming RNA granules after EtBr treatment. U2-OS cells were treated for 5 h with either EtBr or water. Shown are whole cells for *MT-ATP6/8* labeled cells and zoom ins of cells labeled for *MT-ATP6/8*, *MT-CO3* and *MT-ND5*. Arrowheads mark RNA granules. White boxes represent areas that are zoomed in. Scale bars represent 10 μm for whole cells and 2 μm for zoom ins. Cyan represents DAPI staining of nuclei, gray of RNA staining. **(B)** Dynamics of RNA granule formation of *MT-CO3* during EtBr treatment. Cells were treated for 5 hours with water or increasing times with EtBr. Cells were stained with Wheat Germ Agglutinin (WGA) and RNAs were labeled by SABER-FISH. Scale bars represent 5 μm. Arrowheads mark RNA granules. **(C)** Schematic of the fraction of cellular area occupied measurement. Fraction of cellular area occupied is calculated by the RNA area divided by the cell area. A high fraction of cellular area occupied represents a high distribution of mitochondrial RNA in the network, meaning in the cell. A low fraction of cellular area occupied represents a more granular state. **(D)** Quantification of the fraction of cellular area occupied for cells of 3 field of views for each transcript. Single points represent a single cell. Boxplots show the median, the first and third quartiles, the whiskers show a maximum of 1.5 times of the interquartile range. Green represents EtBr treated cells, turquoise represents cells treated with water and the treatment time was 5 hours. The y-axis shows the measured fraction of cellular area occupied in log_10_-scale. n.s. = non significant, **** = p-value > 0.0001. For detailed p-values see also supplemental table 4. **(E)** IMT1B inhibits POLRMT directly. **(F)** Representative images of cells treated for 5 hours either with 10 μM IMT1B or DMSO. Cells were stained with WGA and RNA-FISH was performed for *MT-CO3* and *MT-ND5*. Scale bars represent 5 μm. Arrowheads mark RNA granules. **(G)** Quantification of the fraction of cellular area occupied after IMT1B treatment shown as boxplots. Single points represent a single cell. Green represents IMT1B treated cells, turquoise represents cells treated with water, treatment time was 5 hours. The y-axis shows the measured cellular area occupied in log_10_-scale. **** = p-value > 0.0001, n.s. = non significant. All images in this figure were adjusted to represent the spatial distribution and not differences in intensities.

To confirm that granule formation is directly triggered by transcription arrest, rather than by potential off-target effects of ethidium bromide (EtBr) (King and Attardi, 1989; Toompuu *et al*., 2018), we investigated two additional conditions that inhibit mitochondrial transcription through distinct mechanisms. First, we treated cells with IMT1B, a specific inhibitor of POLRMT (Bonekamp *et al*., 2020). Investigating the spatial distribution of two representative mt-mRNAs, *MT-ND5* and *MT-CO3*, revealed granule formation similar to that observed with EtBr treatment (**Fig. 2E-G**, **EV 2B**). Second, we downregulated SUV3, a subunit of the mitochondrial exonuclease, which leads to an increase of 7S RNA, which promotes POLRMT dimerization and consequently inhibits transcription initiation (Zhu *et al*., 2022). We found that depletion of SUV3 also resulted in the formation of mt-mRNA clusters similar to those observed with drug treatments (**Fig. EV 1F,G**). Collectively, these results demonstrate that inhibition granules form in response to transcription arrest, whether induced pharmacologically or genetically. Thus, the formation of these granules appears to be a consistent cellular response to decreased mitochondrial transcription.

### Inhibition granules are distinct from RNA processing MRGs

We next asked whether the granules described here are distinct from the canonical MRGs. Two RNA-binding proteins, GRSF1 and FASTKD2, are key for the formation of MRGs (Antonicka *et al*., 2013; Jourdain *et al*., 2013; Antonicka and Shoubridge, 2015; Popow *et al*., 2015; Rey *et al*., 2020; Xavier and Martinou, 2021). We used immunofluorescence to examine the localization of GRSF1 and FASTKD2, following treatment with EtBr or IMT1B (**Fig. 3A-D**). Under normal conditions, both proteins displayed characteristic foci within the mitochondrial network, consistent with their known localization in MRGs. However, upon transcription inhibition, neither GRSF1 nor FASTKD2 exhibited the clustering observed for mt-mRNAs. Instead, these MRG markers lost their granular distribution, a finding consistent with previous reports that GRSF1 requires active transcription to form MRGs (Antonicka *et al*., 2013). These observations indicate that the RNA-containing structures formed during transcription inhibition are distinct from canonical MRGs that we therefore designate as “inhibition granules”.

**Figure 3:**
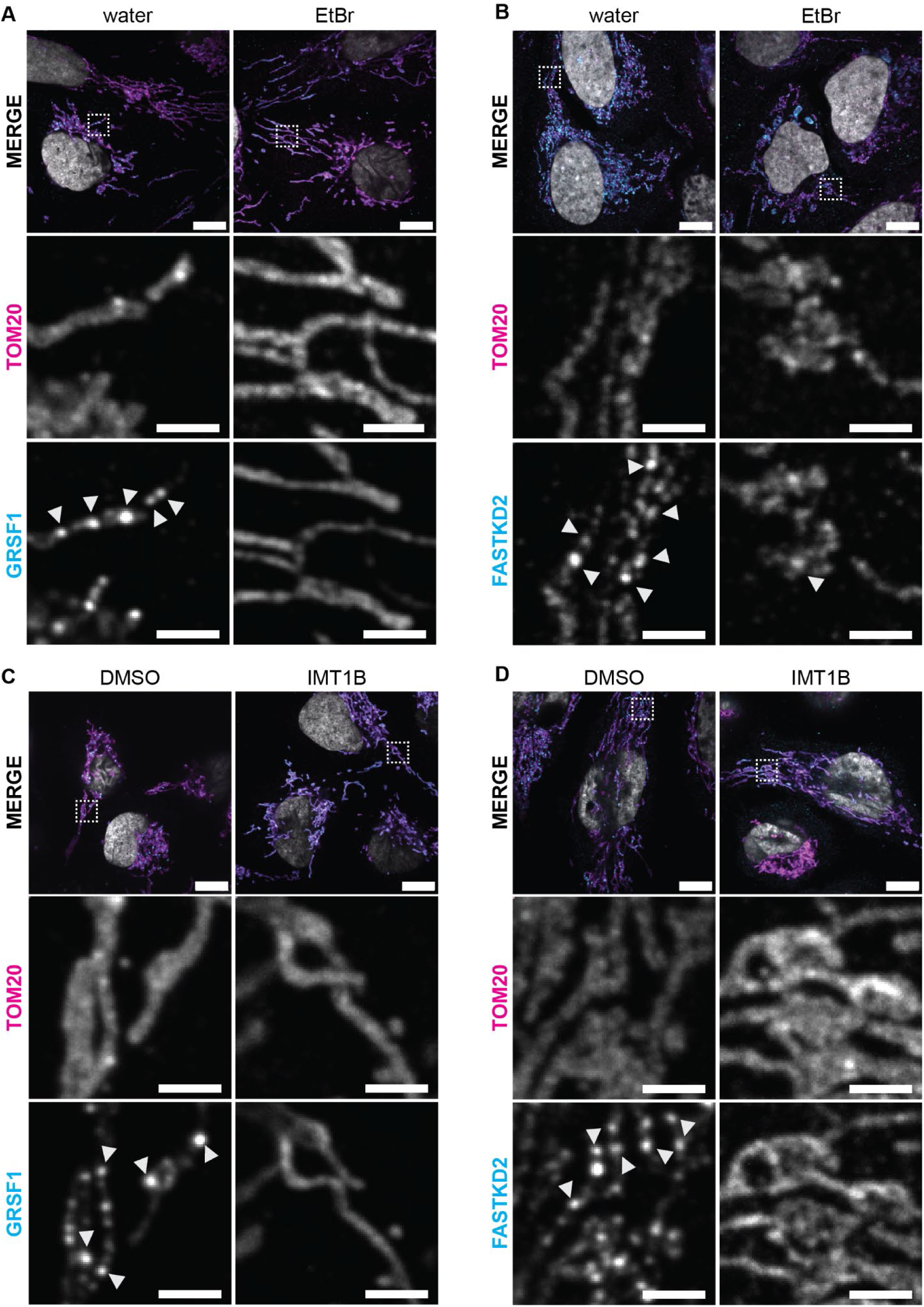
The MRG-markers GRSF1 and FASTKD2 are not part of inhibition granules. (A, **C)** Immunofluorescence of cells treated for 5 hours with EtBr, water, IMT1B or DMSO. Tom20 was used as a mitochondrial marker in all panels. Tom20 is shown in magenta in the MERGE panel and GRSF1 in cyan. **(B, D)** Immunofluorescence of cells treated for 5 hours with EtBr, water, IMT1B or DMSO. Tom20 was used as a mitochondrial marker in all panels.Tom20 is shown in magenta in the MERGE panel and FASTKD2 in cyan. DAPI is shown in gray. Boxes in MERGE panels represent areas enlarged. Zoom ins show RNA staining in gray. Arrowheads mark MRGs. scale bars in whole cell panels represent 10 μm and in zoom-ins 2 μm.

RNA granule formation is influenced by molecular crowding(André and Spruijt, 2020; Hutten and Dormann, 2020) which has been reported to increase in the mitochondrial network upon stress as cristae are disturbed(Bulthuis *et al*., 2023). Hence, a disturbed mitochondrial network could lead to RNA clusters similar to the phenotype we observe. To investigate this possibility, we used mitochondrial markers, the leucine rich pentatricopeptide repeat containing protein LRPPRC or Tom20, to determine the integrity of the mitochondrial network after EtBr or IMT1B treatment (**Fig. 3**, **EV 3A**). In addition, transmission electron microscopy (TEM) was applied to visualize the cristae structure after drug treatment (**Fig. EV 3B**). For EtBr, a few cells showed displacements of cristae and swollen mitochondria (**Fig. EV 3A,B,C**). However, this was not the case for the more selective IMT1B; here, the mitochondrial network was unchanged after 5 hours of treatment (**Fig. 3 C, D**) or even after 24 h of treatment (**Fig. EV 3B**). Thus, the granule formation is not caused by a disturbed mitochondrial network for IMT1B, and could play only a minor role in EtBr.

### Levels of mt-RNAs decrease without changes to nuclear-encoded RNAs

After transcription inhibition, mt-RNAs are expected to decrease with half-lives measured between 20-160 min(Gelfand and Attardi, 1981; Nagao, Hino-Shigi and Suzuki, 2008; McShane *et al*., 2024). By quantifying the overall RNA intensities per cell, we observed a strong decrease for the majority of the tested transcripts in EtBr (**Fig. EV 2C**) or IMT1B treated cells (**Fig. EV 2D**), reflecting a decrease in RNA abundance.

We next asked whether nuclear gene expression changes in response to lowered mt-mRNA levels. RNA-seq analysis after 5 and 24 h of EtBr treatment of HeLa cells showed that mitochondrial-encoded transcripts decrease severely in abundance (**Fig. 4A**), in line with the microscopy data (**Fig. EV 2C**). By contrast, levels of mRNAs of nuclear-encoded OXPHOS subunits did not change (**Fig. 4A**), and neither did other gene groups. Thus, transcription arrest leads to an imbalance in OXPHOS expression, which has been shown to be detrimental to cellular homeostasis(Kruse *et al*., 2008; Kühl *et al*., 2017; Soto *et al*., 2022).

**Figure 4:**
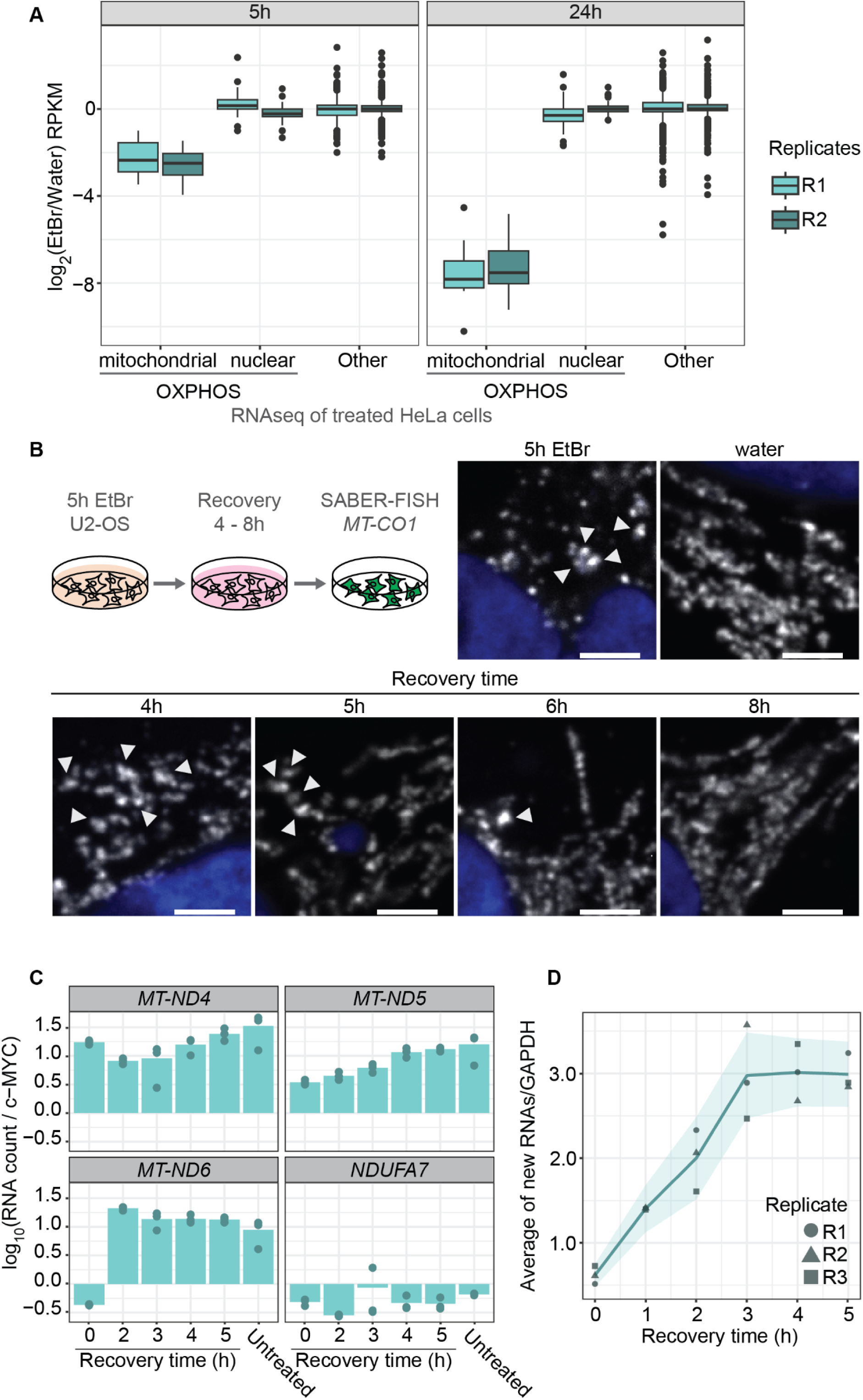
Cells recover from transcription arrest. **(A)** RNA-seq after EtBr or water treatment for 5 or 24 h of HeLa S3 cells. Box plots represent the distribution change for mitochondrial-encoded, nuclear-encoded OXPHOS subunits and other (residual transcriptome) transcripts for two biological replicates. The y-axis shows the log_2_-change of EtBr over the water control in RPKM. **(B)** Confocal microscopy of *MT-CO1* distribution changes in the mitochondrial network during recovery. U2-OS cells were pretreated for 5 hours with EtBr, washed and grown in normal media for 4 up to 8 hours. SABER-FISH for *MT-CO1* was performed and the cells were imaged. Shown is SABER-FISH after 5 h of EtBr and water treatment as controls. Zoom ins are shown for the different recovery time points as well as for the control panels. Whole cells can be found in Fig. EV 4A. Scale bars represent 2 μm. White arrowheads mark RNA granules. All images in this figure were adjusted to represent the spatial distribution and not differences in intensities. **(C)** Relative RNA counts of cells during recovery of IMT1B treated cells. Cells were pretreated with 10 μM IMT1B, washed and grown in normal media. In the last 10 min of each time points 4sU was added. RNA levels were measured after 0, 2, 3, 4, and 5 hours of recovery by MitoStrings and their counts normalized to RNA counts of c-MYC. The y-axis shows the log_10_ scale of relative RNA counts of either *MT-ND4*, *MT-ND5*, *MT-ND6* or *NDUF7A* as a cytosolic control. Shown is the mean of three biological replicates and points represent the individual replicates. **(D)** Meta gene plot of transient transcription analysis during recovery. 4sU labeled RNAs were biotinylated and enriched on streptavidin beads. Eluted RNAs were then measured by MitoStrings. Shown are the averaged relative counts of all 4sU labeled mt-mRNAs normalized to 4sU labeled *GAPDH* on the y-axis. The x-axis shows the time course of 0, 1, 2, 3, 4 and 5 hours. Shown is the mean of the average of three biological replicates in dark green. In dark gray green and in different shapes are the single replicates plotted to show the variation of the experiment. The turquoise ribbon represents the 95% confidence interval.

### Cells resolve inhibition granules upon stress release

We next investigated whether cells can recover and resolve inhibition granules. We treated cells for 5 hours with EtBr and studied RNA distribution during recovery for 4, 5, 6 and 8 h. Interestingly, cells were able to recover as granules disappear after 6 hours of stress release, although they began to dissipate hours earlier (**Fig. 4B** and **EV 4A**).

We next sought to uncover how cells recover from transcription inhibition. We first measured how total mt-mRNA levels change during recovery after IMT1B treatment. RNA levels increased after 5 h of recovery almost to the same amount of untreated cells (**Fig. 4C**), suggesting that transcription restart contributes to the RNA repopulation of the mitochondrial network. To determine the timing of transcription restart, we performed transient transcription analysis during recovery(Schwalb *et al*., 2016). We labeled new RNA with 4sU for 10 min of each time point (0-5 h) and analyzed the fraction labeled in a metagene plot. Already after 1 hour of recovery, there was a slight increase in the newly made fraction of RNAs measured. The amount of newly made transcripts in 10 min increased further overtime, showing that although transcription restarts early, higher rates of transcription emerge slowly (**Fig. 4D**, **EV 4B**).

### Inhibition granules stabilize mt-mRNAs

We reasoned that inhibition granules may form as a reservoir to overcome OXPHOS imbalance during stress recovery. Consistent with granules serving a protective role, RNAs remained visible after 24 h of drug treatments, despite the short half-lives of mt-mRNA at steady-state (**Fig. 5A**, **Fig. EV 1E**). To obtain a quantitative view of mt-mRNA decay after transcription inhibition, we measured RNA abundance through time after transcription inhibition by either EtBr or IMT1B to measure half-lifes at early and late timepoints. If inhibition granules are indeed protective, stabilization would be observed in the kinetics of RNA decay as a 2-state model, where one population (or state) is degraded fast and one stays more stable. For cells treated with EtBr, we detected a two-state decay curve for *MT-ND5* and *MT-ND6*. In contrast, the nuclear-encoded control transcript *NDUF7A* had a decay curve with a one-state model (**Fig. 5B** and **Fig EV 5A**, **Table 1**). For cells treated with IMT1B treatment, we were able to detect more transcripts with decay curves consistent with a 2-state model (*MT-ND1, MT-ND2, MT-ND3, MT-ND5* and *MT-ND6*) (**Fig. 5C** and **Fig. EV 5B**). Thus, inhibition granules are protective for at least some transcripts.

**Figure 5:**
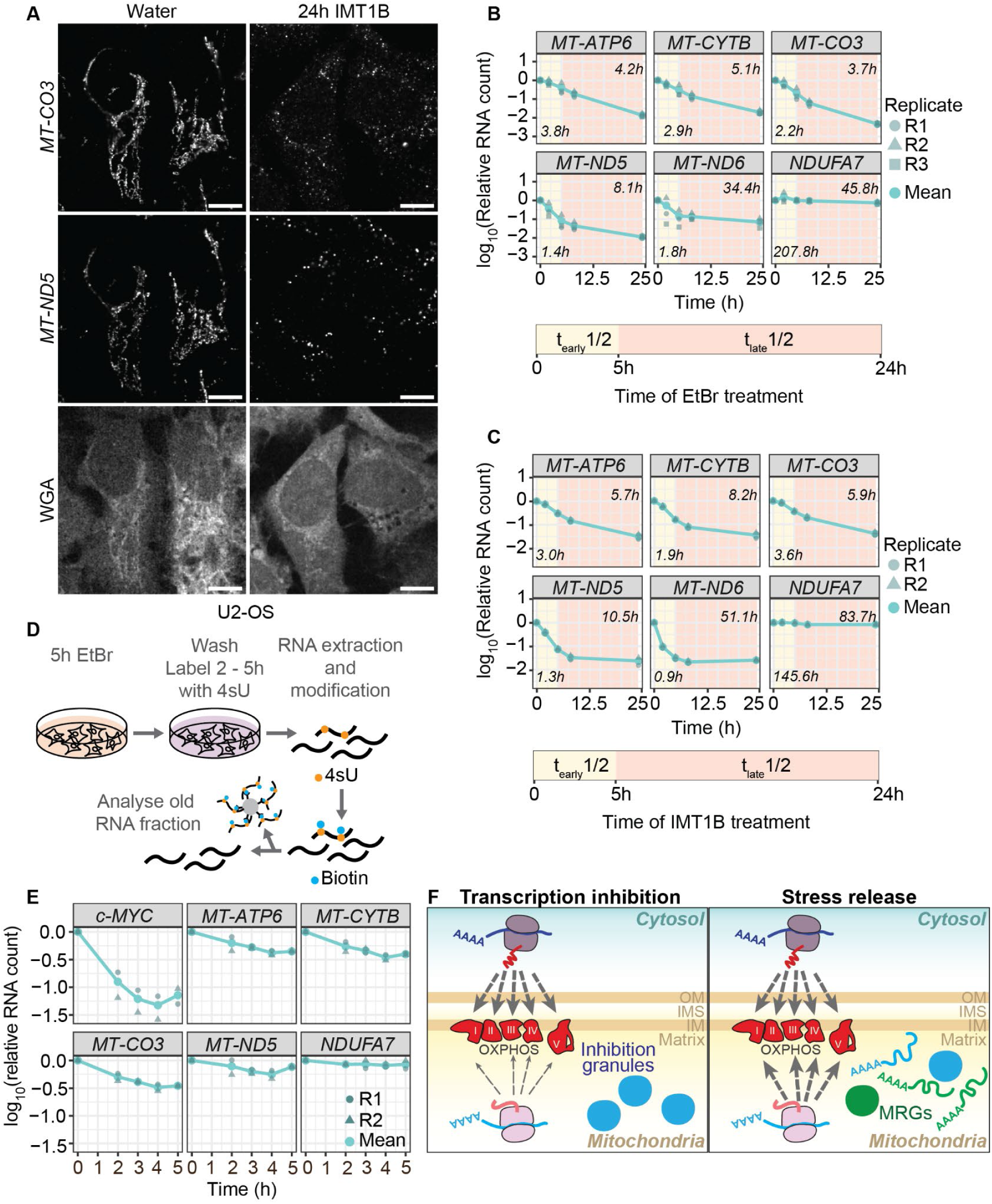
Inhibition granules show protectivity for parts of mt-mRNAs. **(A)** SABER-FISH for *MT-CO3* and *MT-ND5* after 24 h of IMT1B or DMSO treatment in U2-OS cells. RNAs are shown in gray as well as WGA staining of cells. scale bars represent 10 μm. Signals were adjusted to represent the spatial distribution and not for intensity comparisons. **(B)** Degradation kinetics of representative mt-mRNAs in EtBr long-term treatment. Cells were incubated for 0 up to 24 h in EtBr media. RNAs were extracted and RNA abundance was measured by MitoStrings. The y-axis shows the log_10_ of the RNA counts normalized to the counts of *GAPDH*. The x-axis shows the treatment time in hours. The bar underneath represents the time frames used to calculate half-lifes for early (t_early_1/2) and late time points (t_late_1/2). Shown is the mean of three biological replicates in turquoise. In dark green and different shapes the single biological replicates are shown. The numbers in the left lower corner represent early half-lifes and in the upper right corner the late half-lifes. **(C)** Degradation kinetics of representative mt-mRNAs in IMT1B long-term treatment. Cells were incubated for 0 up to 24 h in IMT1B media. RNAs were extracted and RNA abundance was measured by MitoStrings. The y-axis shows the log_10_ of the RNA counts normalized to the counts of *GAPDH*. The x-axis shows the treatment time in hours. The bar underneath represents the time frames used to calculate half-lifes for early (t_early_1/2) and late time points (t_late_1/2). Shown is the mean of two biological replicates in turquoise. In dark green and different shapes the single biological replicates are shown. The numbers in the left lower corner represent early half-lifes and in the upper right corner the late half-lifes. **(D)** MitoStrings analysis of representative unlabeled granular RNAs after stress release. Cells were treated for 5 hours with EtBr, washed and grown in media supplemented with 4sU for 0, 2, 3, 4 and 5 hours. RNAs were extracted and 4sU (shown in orange) labeled RNA was biotinylated (biotin is shown in blue). Biotinylated RNA was bound onto streptavidin beads and the flowthrough, representing the unlabeled RNA fraction, was used for MitoStrings. **(E)** Graph showing decrease of unlabeled RNAs after stress release. The y-axis shows the log_10_ of RNA levels normalized to an unlabeled spike-in, added during RNA extraction. **(F)** During transcription inhibition mitochondrial RNAs form inhibition granules and the OXPHOS expression inside mitochondria is decreased whereas nuclear-expression of OXPHOS subunits remains unchanged. During stress release OXPHOS expression will eventually increase again. Inhibition granules might help in recovery through release of stored RNAs. MRGs will form again and replenish the mitochondrial network with RNAs. Inhibition granules are shown in blue, MRGs in green.

**Table 1:**
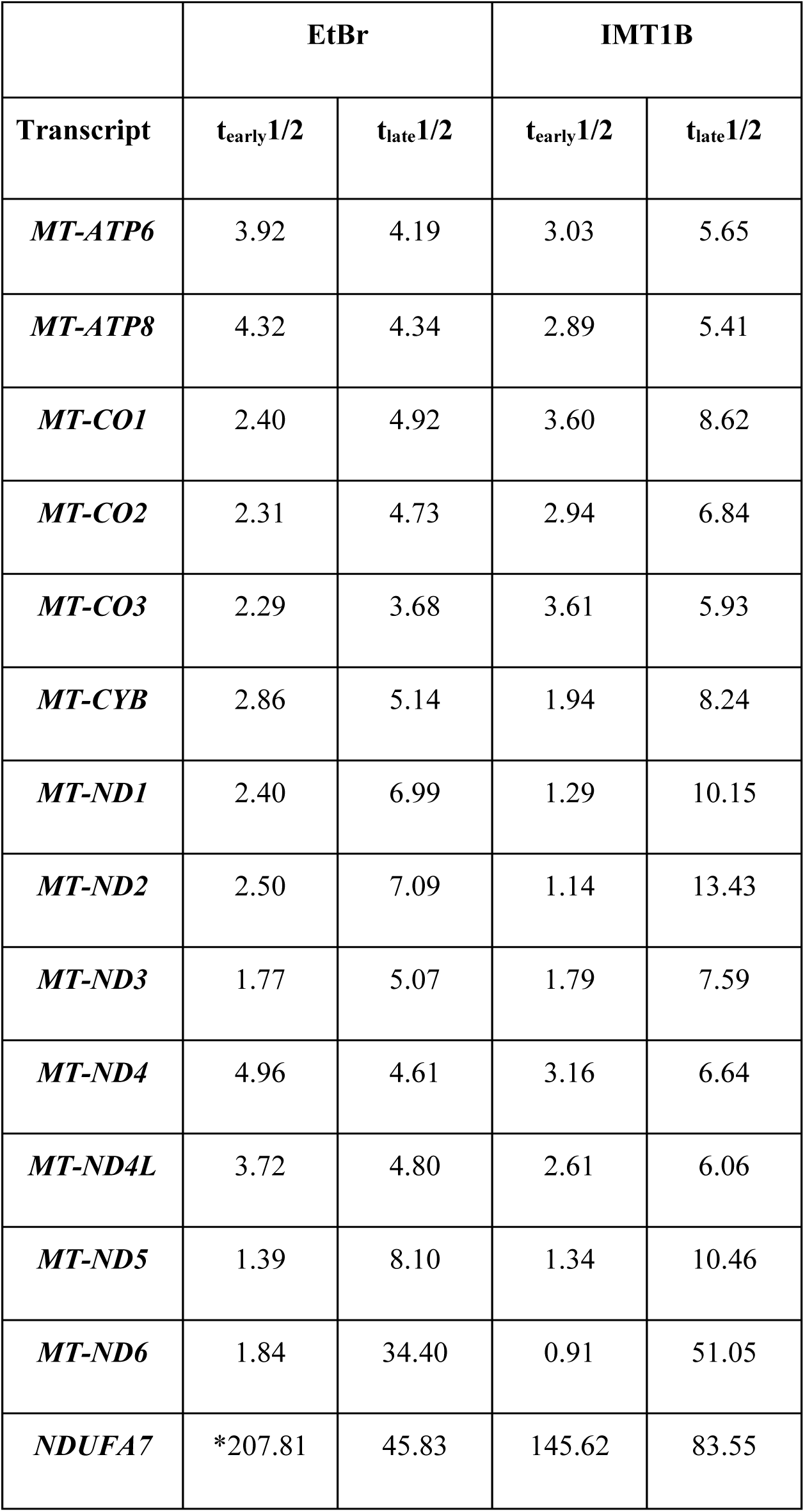
Half-lifes of long-term treated RNAs. *=only 0 and 5 hours were used for linear regression as the half-life resulted otherwise in negative values.

To elucidate the fate of RNAs within inhibition granules during stress release, we employed a metabolic labeling approach. After treating cells with IMT1B for 5 hours to induce inhibition granules, we removed the inhibitor and immediately added 4sU to label all newly synthesized RNAs (**Fig. 5D,E**). This strategy allowed us to distinguish between pre-existing and newly transcribed RNAs. We then measured the levels of unlabeled RNAs, which were predominantly those that had been sequestered in inhibition granules. Our analysis revealed a gradual decrease in the abundance of these pre-existing RNAs, beginning approximately 2 hours after stress release (**Fig. 5D,E** and **Fig. EV 5C**). This decline, although slow, indicates that the RNAs are being degraded. The onset of RNA degradation coincides with the timing of inhibition granule dissolution, suggesting that mt-mRNAs lose their protective environment as the granules dissipate, becoming susceptible to degradation processes.

In this study, we have identified a novel type of mitochondrial RNA granule that forms in response to transcription inhibition. Our experiments employed strong perturbations to induce widespread transcription arrest across the mitochondrial network, resulting in the formation of large, easily visualized granules that persisted for hours. However, we hypothesize that smaller, transient granules may form under physiological conditions as individual mtDNA molecules switch between transcriptionally active and inactive states. During transcription arrest, we observed an imbalance in OXPHOS expression, coinciding with the sequestration of RNAs into these inhibition granules. We propose that these structures act as RNA reservoirs, facilitating recovery after stress release or upon transcription reactivation (**Fig. 5F**).

The formation of these granules likely involves various molecular interactions, including protein-protein, RNA-protein, and RNA-RNA associations (Van Treeck *et al*., 2018; Matheny *et al*., 2021). Mitochondrial RNA-binding proteins, particularly members of the FASTK family known for their roles in RNA stability and processing, are potential candidates for mediating inhibition granule assembly (Jourdain *et al*., 2015; Popow *et al*., 2015; Boehm *et al*., 2017; Ohkubo *et al*., 2021). The discovery of inhibition granules adds to the growing list of mitochondrial RNA granules and underscores the complexity of mitochondrial gene expression regulation. These findings reveal an additional layer of compartmentalization within mitochondrial gene expression, that together with other mitochondrial RNA granules provide multiple intervention points throughout the gene expression process, allowing cells to maintain proteostasis in the face of aberrant OXPHOS expression.

## Acknowledgements

We thank the Churchman lab for helpful discussion. We thank R. Stefan Isaac for helpful discussions and comments. We thank Nick Kramer for help with imaging. We are grateful to the MicRon facility at HMS, especially Paula Montero Llopis and Praju Vikas Anekal for advice and help with imaging and analysis. We want to thank the Biopolymers Core Facility at Harvard Medical School for sequencing services and the Boston Children’s Hospital Molecular Genetics Core, the Bauer facility at Harvard and the Manning lab at Harvard T.H. Chan School of Public Health for NanoStrings services. **Funding:** This work was supported by the NIH (R01-GM123002 to L.S.C.), a Helen Hay Whitney Foundation Fellowship (F-1240 to K.G.H.), EMBO fellowship (ALTF 762-2019 to K.G.H.) and PROMOS fellowship to C.A.M.W.. **Author Contributions:** K.G.H., E.M., C.A.M.W. and L.S.C. designed the experiments. K.G.H., C.A.M.W., A.K.B. and E.M. performed the experiments. K.G.H. performed the computational analyses. K.G.H. and L.S.C. wrote the manuscript. **Competing interests:** The authors declare no competing interests.

---

**Figure EV1:**
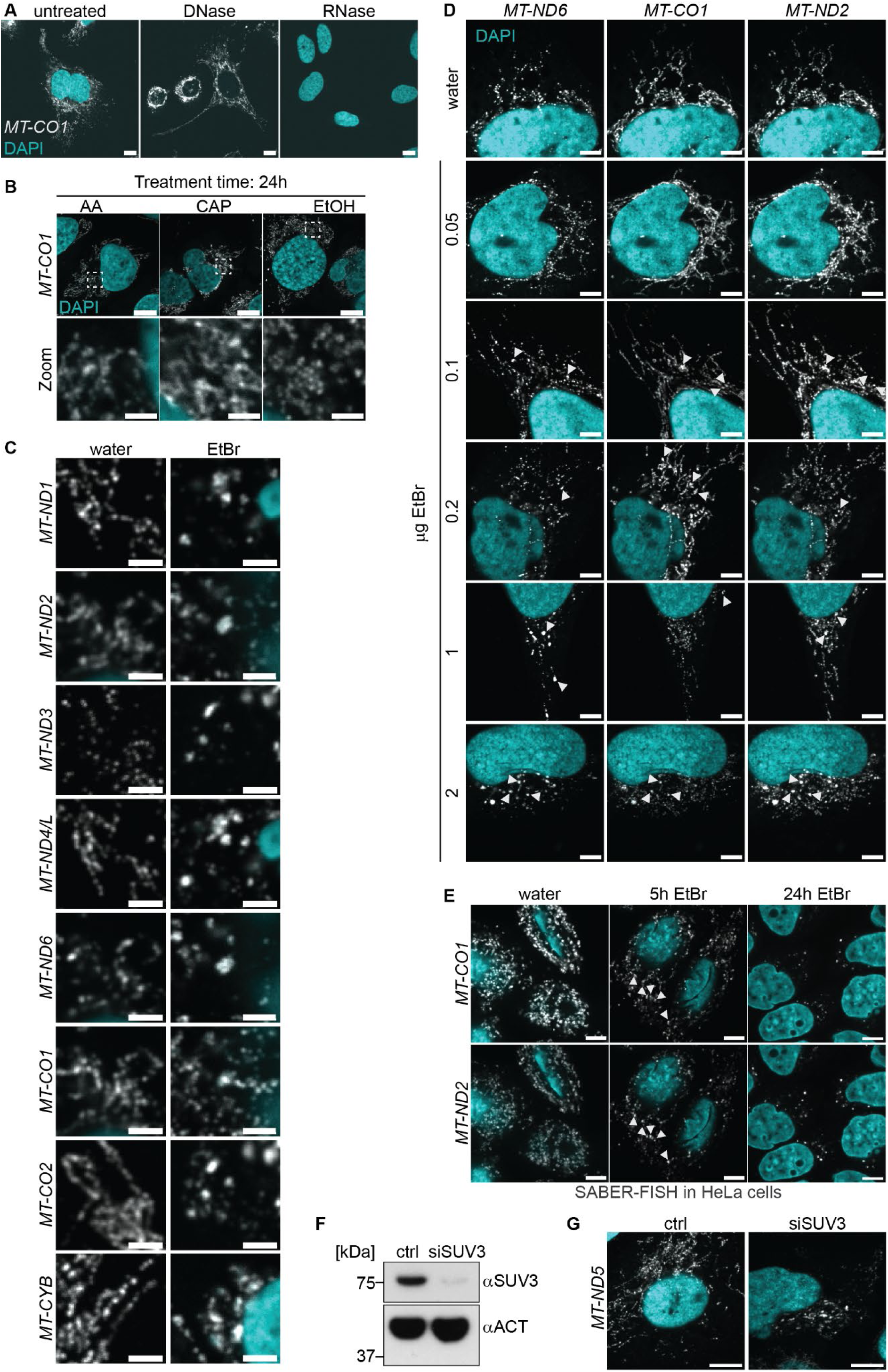
Mitochondrial RNAs form granules upon EtBr treatment. **(A)** RNAse and DNAse treatment to test for oligo specificity of *MT-CO1*. Cells were either treated after fixation and before the hybridization. Scale bars represent 10 μm. **(B)** SABER-FISH for *MT-CO1* after 24 hours of drug treatment with 100 nM antimycin A (AA), 200 μg/ml chloramphenicol (CAP) or EtOH as a control. DAPI stain is shown in cyan, RNAs in gray. White boxes mark areas chosen for zoom-ins. Scale bars of whole cell panels represent 10 μm and in zoom-ins 2 μm. **(C)** Example images for residual mt-mRNAs not shown in Fig. 2A. All mt-mRNAs form granules upon EtBr treatment. Scale bars represent 2 μm. **(D)** Titration of EtBr treatments for 5 h. First granules can be detected visually with 100 ng/ml EtBr. Shown are representative images of multiplexed *MT-ND6*, *MT-ND2* and *MT-CO1*. RNAs are presented in gray, DAPI staining of nuclei in cyan. Arrowheads mark RNA granules. Scale bars are 5 μm. **(E)** SABER-FISH of EtBr treated HeLa cells. RNAs are presented in gray, DAPI staining of nuclei in cyan. Arrowheads mark RNA granules. Scale bars are 5 μm. **(F)** Control western blot of U2-OS cells treated with a non-targeting siRNA (ctrl) and siRNAs targeting SUV3 for 72 hours. SUV3 depletion was highly effective. **(G)** SABER-FISH after siRNA treatment to deplete SUV3. Cells were treated for 72 h. RNAs are presented in gray, DAPI staining of nuclei in cyan. Scale bars are 10 μm. All images in this figure were adjusted to represent the spatial distribution and not differences in intensities.

**Figure EV2:**
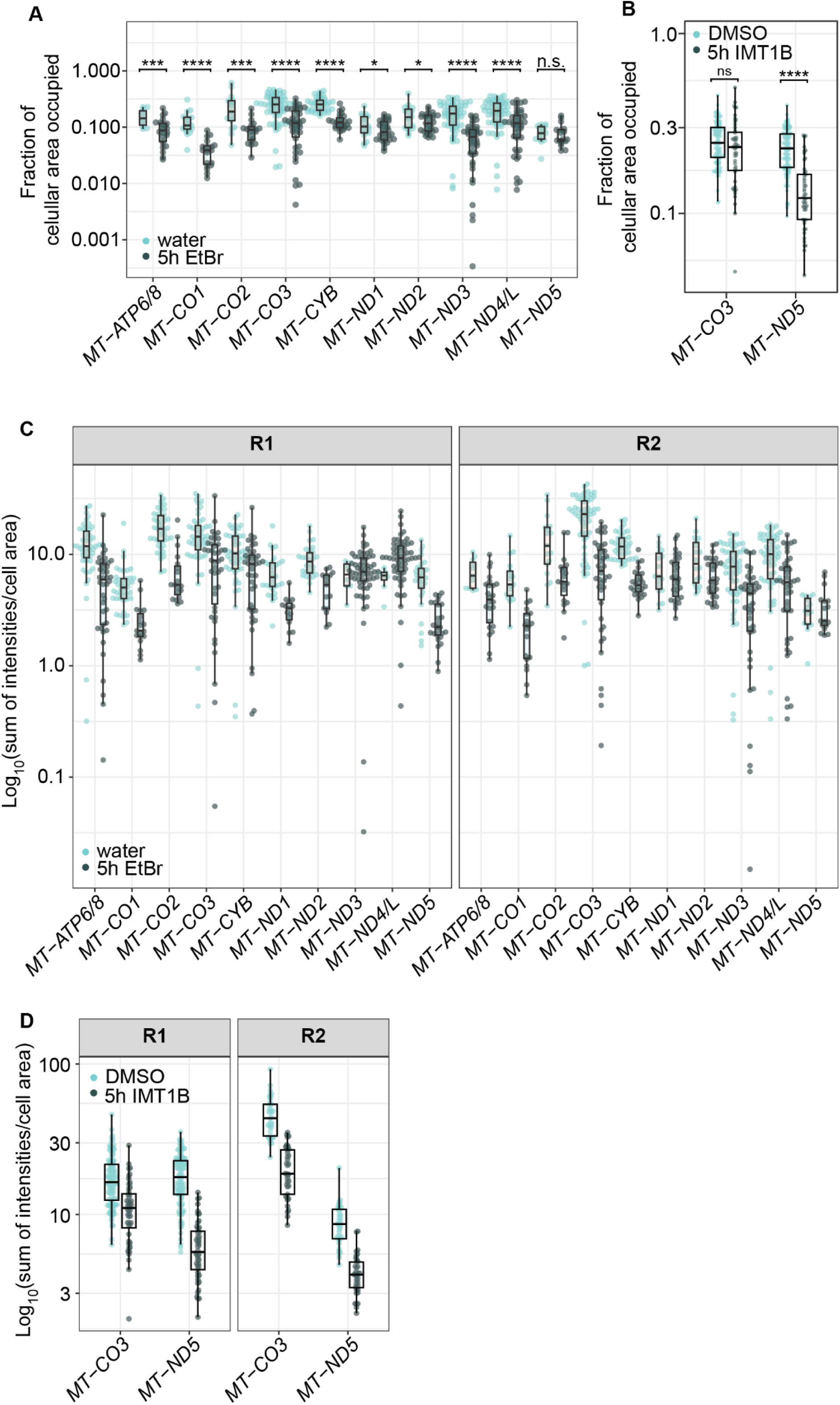
RNA abundance decreases upon transcription arrest. (A, **B)** Single cell data of biological replicates of quantified fraction of cellular area occupied after either EtBr or IMT1B treatment. Single points represent a single cell. Boxplots show the median, the first and third quartiles, the whiskers show a maximum of 1.5 times of the interquartile range. Green represents drug treated cells, turquoise represents cells treated with water or DMSO, treatment time was 5 hours. The y-axis shows the measured cellular area occupied in log_10_-scale. n.s. = non significant, * = p-value > 0.05, *** = p-value > 0.001, **** = p-value > 0.0001. For detailed p-values see supplemental table 4. **(C)** Boxplots showing the distribution of intensities measured in cells. Single points represent a cell. To measure intensities, the integrated intensities measured of RNA objects were summed per cell and normalized to the cell area. The y-axis presents the normalized summed intensities on a log_10_ scale. Drug is shown in dark green, the vehicle control in turquoise. **(D)** Intensity measurements of single cells upon IMT1B treatment. Intensities are represented as described in panel (C).

**Figure EV3:**
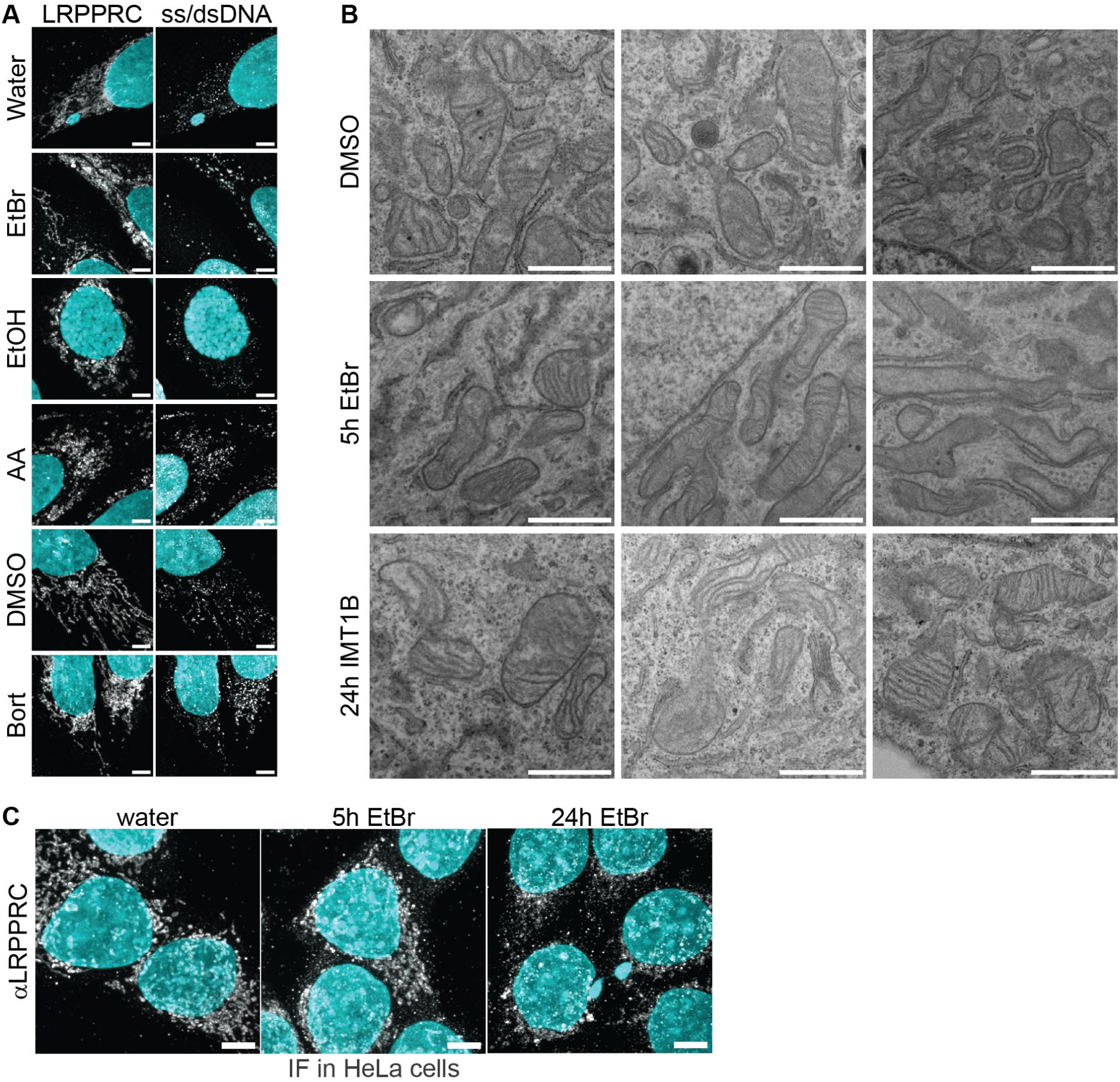
The integrity of the mitochondrial network is not strongly affected. **(A)** Immunofluorescence for LRPPRC and mitochondrial DNA (ss/dsDNA) after different drug treatments is shown. Cells were treated with Bort (bortezomib), AA (antimycin A) or EtBr (ethidium bromide) and the respective vehicle controls. LRPPRC and IF labeled DNA are shown in gray. DAPI stain is shown in cyan. Some cells show swollen mitochondria after EtBr treatment whereas others have normal mitochondria. Scale bars show 5 μm. All images in this figure were adjusted to represent the spatial distribution and not differences in intensities. **(B)** Transmission electron microscopy (TEM) of cells treated for 24 h with IMT1B, DMSO or for 5 h with EtBr. Bars show 1 μm. Shown are three zooms of three different cells. Mitochondrial morphology is not majorly disturbed. **(C)** Immunofluorescence for LRPPRC in HeLa cells after EtBr treatment for 5 or 24 hours. LRPPRC is presented in gray, whereas cyan shows the nuclei stained by DAPI. The mitochondrial network is affected after 24 hours of EtBr treatment. Scale bars show 5 μm. All images in this figure were adjusted to represent the spatial distribution and not differences in intensities.

**Figure EV4:**
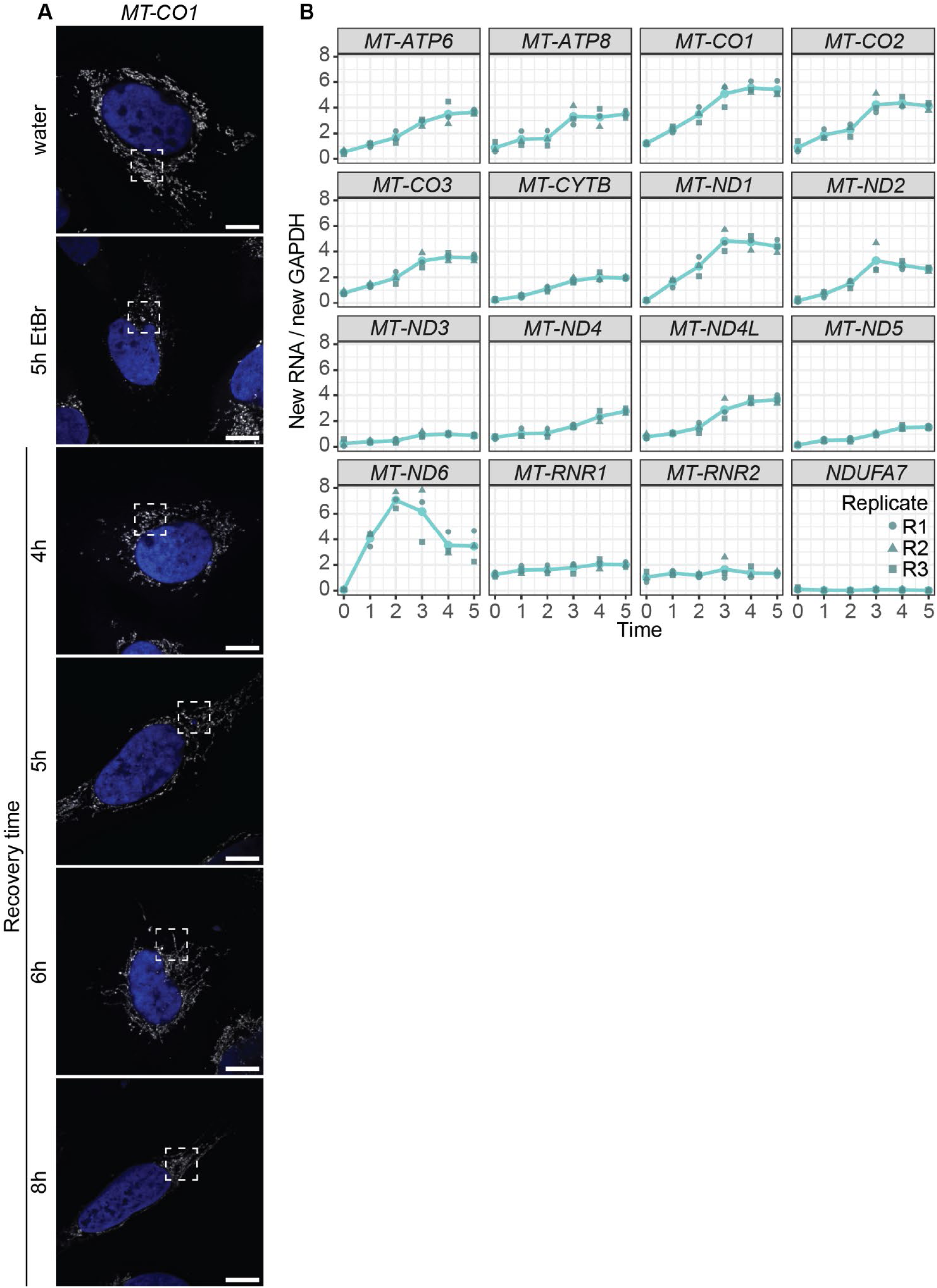
Cells recover after release of transcription arrest. **(A)** Whole cell images corresponding to zoom-ins shown in Fig. 4C. *MT-CO1* is shown in gray and DAPI in blue. White boxes mark areas used for zoom-ins. Scale bars represent 10 μm. Images are adjusted to show the spatial distribution of RNAs and not to compare intensities. **(B)** Transient transcription during recovery after IMT1B treatment. Shown are relative RNA counts of 4sU labeled RNAs normalized to 4sU labeled GAPDH on the y-axis. The x-axis shows the time points of recovery in hours. In turquoise the mean of three biological replicates is shown as a line plot. The biological replicates are shown in dark green and in different shapes.

**Figure EV5:**
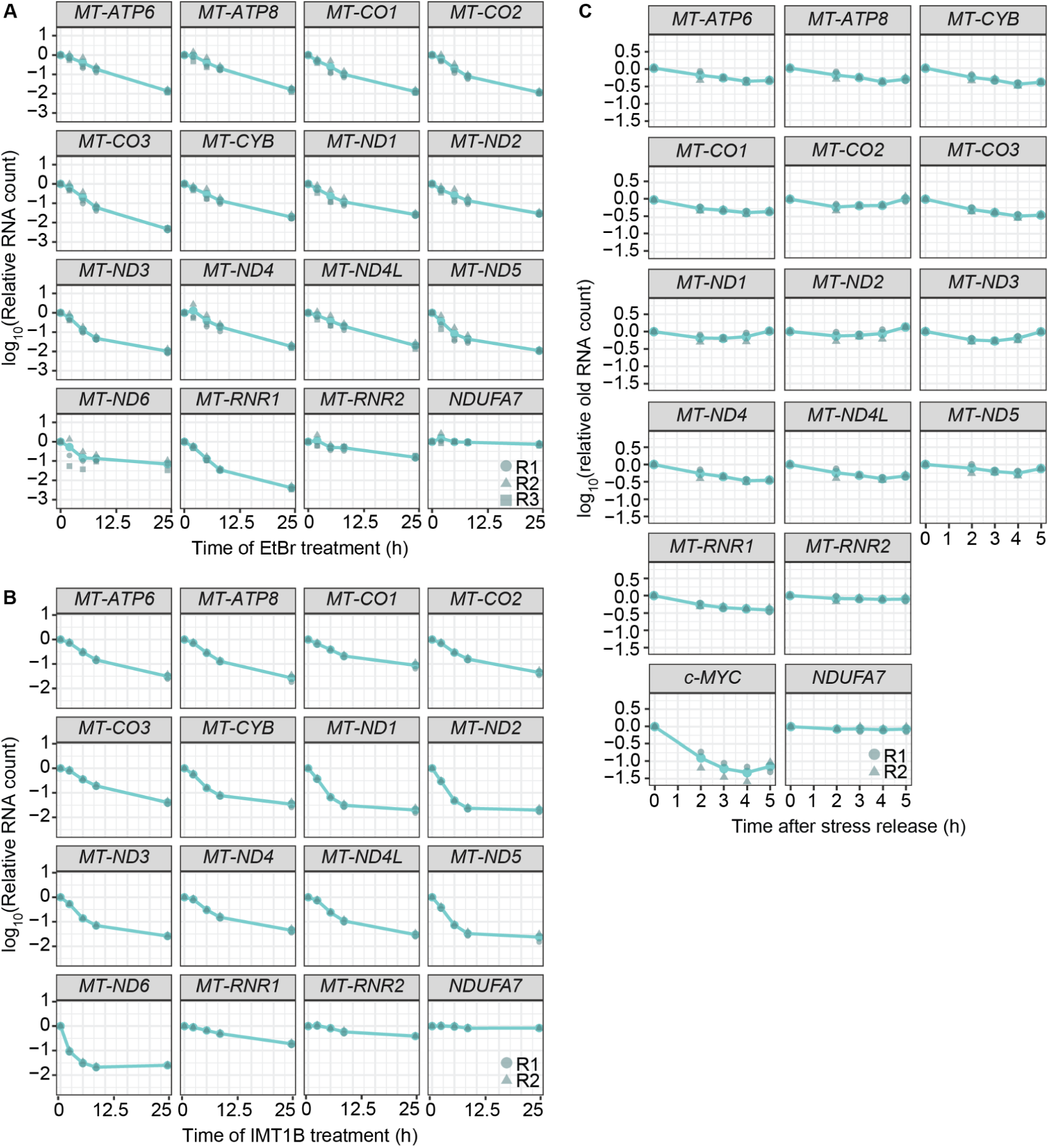
Degradation kinetics after long-term stress treatments. **(A)** Shown are the different degradation kinetics of the different mitochondrial RNAs during long-term EtBr treatment. Presented is the mean of three biological replicates. Single replicates are shown in dark green and as different shapes. The y-axis shows log_10_ of RNA counts normalized to *GAPDH* counts. Single transcripts are shown as representatives in Fig. 5B. **(B)** Graphs represent the different degradation kinetics of the different mitochondrial RNAs during long-term IMT1B treatment. Shown is the mean of two biological replicates as well as the single replicates in dark green and as different shapes. Representative transcripts of this set are shown in Fig. 5C. **(C)** Shown is the change of old RNA levels upon stress release. The mean of two biological replicates is shown in turquoise and individual replicates in dark green. The RNA counts were normalized to an unlabeled spike-in. The y-axis shows the log_10_ of spike-in normalized RNA counts. Representative transcripts of this set are shown in Fig. 5E.

## Material and methods

### Cell culture

U2-OS cells (ATCC) were cultured in McCoy’s 5A media (ATCC #30-2007) supplemented with 10% FBS. HeLa cells were cultured in DMEM media supplemented with 10% FBS. For cell passaging, cells were rinsed with 1x PBS and treated with Gibco™ TrypLE™ Express Enzym (Gibco™ 12605010) for 15 min (U2-OS cells) or 5 min (HeLa cells). Cells were seeded onto plates with fresh media.

### Drug treatments of cells

Transcription was inhibited either with 2 µg/ml Ethidium bromide or 10 µM IMT1B. The drugs were diluted to the respective concentration in medium and added to the cells for the indicated time points. As controls served 1% water for the ethidium bromide treatment and 0.1% DMSO for IMT1B. 100 µM antimycin A in EtOH or 100 µM bortezomib in DMSO were used in Fig. 1D. As controls, cells were treated with 2% EtOH or 2% DMSO. Treatment time was 5 hours. For treatments of cells with 100 nM antimycin A or 200 µg/ml chloramphenicol for 24 hours, 0.001% EtOH and 0.04% EtOH served as control respectively (Fig. EV 1B).

### siRNA treatment against SUV3

siGENOME Human SUPV3L1 (SUV3) siRNA was ordered in SMARTPool format from Horizon. For siRNA treatment, cells were grown until 70% confluency. Afterwards, they were transfected using the Liptofectamine^TM^ RNAiMAX kit (Invitrogen #13778075). Therefore, 9 μl Liptofectamin was premixed with 350 µl Opti-MEM^TM^ medium (Gibco^TM^ #31985062). 1.5 µl of 20 µM siRNA pools (siSUV3 and untargeted as a control) were mixed with 350 µl Opti-MEM^TM^. Both mixtures were combined and incubated for 5 min. In the meantime, cells were supplemented with fresh media. 250 µl of the mixed siRNAs were added to the respective wells. After 24 hours of siRNA treatment, the media was replaced with fresh media lacking siRNA. After an additional 48 hours, cells were harvested. Some cells were seeded into glass chambers to grow overnight and could be used for SABER-FISH. A fraction of cells was frozen down to be used for western blot analysis.

### Probe design

RNA-FISH probes were designed according to Beliveau et al.(Beliveau *et al*., 2018). Since the mitochondrial genes overlap with pseudogenes, the probe candidates were aligned against the mitochondrial DNA alone. If the whole genome was used no specific probes could be selected, due to pseudogenes in the nuclear DNA. No kmer filtering was used. Where necessary, probe sequences were converted to the reverse complement sequence.

After Oligominer filtered suitable oligo sequences, they were manually extended by primer sequences to be able to do nucleotide exchange reactions with hair pins to synthesize the final oligos. All oligo sequences are listed in supplemental table 1. Hairpin sequences were taken from Kishi et al.(Kishi *et al*., 2019). For bi-cistronic transcripts (*MT-ATP6/8*, *MT-ND4/L*) probe pools were designed that detect the full bi-cistronic transcripts.

### Probe synthesis

Probe synthesis was performed as published (Kishi *et al*., 2019). A reaction mix without oligos was prepared on ice and incubated for 15 min at 37°C to remove any free guanines in the mix. One reaction mix contained 0.5 μM (HP27 and HP28) to 1 μM hair pins (all other hair pins), 1x PBS buffer, 300 nM dNTPs without guanines, 100 nM Clean.G oligo, 10 mM MgSO_4_ and 8U of Bst LF polymerase (BioVision #M1213-200) or 100U Bst LF polymerase (McLab #BPL-200). After the 15 min preincubation 1 µM of oligo pool mix was added. The reaction was incubated for 80 min at 37°C followed by 20 min at 80°C to inactivate the polymerase and cooled down to 4°C. Next, the elongation was tested on 1.3% agarose gels. If an elongation of at least 100 bp was observed, probes were purified using the MINelute PCR reaction kit from Quiagen. The ssDNA probes were eluted using 30 µl of RNAse-free water and their concentration measured by Nanodrop.

### Coating of slides

150 µl of poly-L-lysine was added to each well and incubated for 10 min at RT. After three rinses with 1x PBS, slides were left to dry for 1 hour and could be used for cell seeding.

### SABER-FISH

The SABER-FISH protocol was adapted from Kishi et al.(Kishi *et al*., 2019). Cells were grown in poly-L-lysine coated ibidi glass bottom well chambers (ibidi, #80807) until 50-70% confluency. For drug treatments, cells were grown overnight in normal media and at the next day treated as described in the drug treatment section. After the treatment cells were rinsed 2x with 1x PBS and fixed with 4% freshly prepared formaldehyde (Pierce™ 16 % Formaldehyde (w/v), methanol free, Sigma, #28906) for 1 hour at RT. Fixation time for Fig. 1B, D, Fig. EV1A, C, E, Fig. EV 3A and C has been 20 min. After fixation, cells were rinsed 3x with 1x PBS. Cells would be either stored at 4°C overnight to be processed the next day or directly subjected to permeabilization. In case of SABER-FISH data used for quantification, cells were subjected to WGA staining right before permeabilization. Therefore, cells were treated with 5 µg/ml WGA-CF®405S (Biotium, #29027-1) for 30 min in the dark at RT. The cells were washed three times with 1xPBS and could be permeabilized afterwards. To this end, cells were incubated in 1x PBS and 0.1% TX-100 for 20 min at RT. Afterwards, cells were rinsed 2x with 1x PBS, followed by a 5 min wash in 1x PBS at RT. A wash for 5 min in 1x PBS and 0.1% Tween-20 followed. Cells were equilibrated for 15 min at 43°C in prewarmed Whyb LD buffer (2xSSC, 40% formamide, 0.1% Tween-20). During this incubation the hybridization mix was prepared. 1 µg of each probe was mixed with the 1.4x hybridization buffer (2.8x SSC, 56% Formamide, 0.14 % Tween-20, 14% Dextran). The ratio of hybridization buffer and water including probes was 85 µl to 35 µl per well. The mix was prewarmed to 43°C and the Whyb was replaced with 120 µl of the hybridization mix. The cells were incubated at 43°C overnight. The next day, cells were first quickly rinsed with prewarmed Whyb and washed two times for 30 min at 43°C with Whyb buffer. Two washes at 43°C for 5 min with 2x SSC and 0.1% Tween-20 followed. Cells were rinsed for 1 min with 1x PBS buffer at RT. They were kept in fresh 1x PBS buffer until the hybridization oven cooled down to 37°C. In the meantime the imager probes were mixed for fluorescence detection. From here all incubations were done in the dark to avoid bleaching of the fluorophores. The imager mix contained 1 µM of each imager probe and 1x PBS. Cells were incubated either for 30 or 60 min with imagers at 37°C. The cells were washed with prewarmed 1x PBS once for 5 min and two times for 2 min. Except for WGA stained samples, DAPI staining followed. Therefore, cells were rinsed once with 1x PBS at RT and incubated for 10 min in DAPI (5 µg/ml in 1xPBS and 20 U/µl of SUPER RNAsin) at RT and washed 3 times for 5 min with 1x PBS. 150 µl mounting media (80% glycerol, 1xPBS, 20 mM Tris pH 8, 2.5 mg/ml n-Propyl-Gallate, 20 U/µl SUPER RNAsin) was added and the cells were imaged either the same or next day. For Fig. 1A, D, Fig. EV1A, C and E, the Hyb buffer and Whyb contained 1% Tween-20.

### Immunofluorescence

For immunofluorescence after SABER-FISH treatments, mounting media was removed by two quick rinses with PBST buffer (1xPBS, 0.1% TX100). Cells were washed 3 to 4 times for 5 min with PBST followed by three washes of 5 min with displacement buffer (50% formamide, 1xPBS) to remove fluorescent probes. Next, cells were washed three times for 2 min with PBST and rinsed three times with 1xPBS. Cells were blocked for 1 hour at room temperature with blocking buffer (5% goat serum in 1xPBS and 0.1% Tx100). Cells were incubated overnight at 4°C with the primary antibodies.

Unbound antibodies were removed by three washes for 5 min with PBST at RT. Cells were incubated with a secondary antibody harboring the fluorophores diluted in blocking buffer for 1 hour at RT and washed again three times for 5 min with PBST at RT. After staining nuclei by DAPI, cells were covered with mounting media and were stored at 4°C until imaging.

For immunofluorescence directly after drug treatments, cells were fixed and permeabilized as described in the SABER-FISH section. Afterwards, cells were incubated with blocking buffer and treated as outlined above.

### Antibodies

Listed are antibodies used in this study with information of dilution and where they are purchased from. All antibodies were diluted in blocking buffer, if not stated otherwise: TOM20 (Santa Cruz, #sc-17764) used in a 1:200 to 1:500 dilution, GRSF1 (Abcam, #ab205531) used in a 1:500 to 1:1000 dilution, FASTKD2 (ProteinTech, #17464-1-AP) used in a 1:500 dilution, ds/ssDNA (PROGEN, #61014) used in a 1:100 dilution, LRPPRC/GP130 (Abcam, #ab97505) in a 1:200 dilution, SUV3L (Abcam, #ab127909) used in a 1:1000 dilution in milk for western blot, ACTB (Cell Signaling, #3700) in a 1:5000 dilution in milk for western blots, horseradish peroxidase (HRP)-conjugated anti-rabbit IgG (Cell Signaling, cat. no. 7074S) 1:10000 diluted in 5% milk, HRP-conjugated anti-mouse IgG (Cell Signaling, cat. no. 70765) 1:10000 diluted in 5% milk, goat anti-rabbit IgG Alexa Fluor 647 (Thermo Fisher, cat. no. A-21245) in a 1:1000 dilution, goat anti-mouse IgG2a Alexa Fluor 488 (Thermo Fisher, cat. no. A-21131) in a 1:1000 dilution and goat anti-mouse IgM Alexa Fluor 555 (Thermo Fisher, cat. no. A-21426) in a 1:1000 dilution.

### Western blotting

Cells were lysed in RIPA buffer and treated with Benzonase for 30 min and afterwards mixed with loading buffer (95 mM Tris–HCl, pH:6.8, 7.5% glycerol, 2% SDS, 0.5 mg/ml bromophenol blue, 50 mM DTT). Proteins were separated on 10% NuPAGE Novex Bis-Tris gels (Thermo Fisher Scientific) and transferred to a nitrocellulose membrane. Membranes were incubated with primary antibodies overnight (SUV3L and ACTB). For visualization, membranes were incubated with secondary antibodies conjugated with horseradish peroxidase for chemiluminescence detection.

### Imaging

Samples were imaged on a Nikon Ti inverted microscope with a W1 Yokogawa Spinning disk with 50 µm pinhole disk, a Nikon motorized stage, a Physik Instrument Piezo Z motor, an Andor Zyla 4.2 Plus sCMOS monochrome camera VSC-06522 or VSC-10150 (used for quantification images of replicates 2 of IMT1B treatment and *MT-ND5* of EtBr treatment) and OKO Lab Heated enclosure with CO_2_ control at room temperature. For imaging the Nikon Elements Acquisition Software AR 5.02 was used. All files were saved in .ND2 format. Before each imaging, slides were cleaned first with Sparkle and water to remove salts and finally with 70% EtOH to remove any oils. Samples for quantifications were imaged with a Plan Apo λ 60x/1.4 Oil DIC objective in combination with immersion oil (Cargille, non-drying immersion oil Type 37, #16237) and with a laser device of Digital Mirror Device lines (Aura/LED), 350, 380, 488 and 561 with emission filter 455/50, 480/40, 525/36, 605/52, 630/75 and 705/72 or with laser lines 405, 488, 561, 640 (Nikon Instruments, Laser Unit model Lun-F) and emission filters (455/50, 480/40, 525/36, 605/52, 630/75, 705/72). For each image set, 10 field of views of an empty ibidi glass chamber well were imaged with the same laser settings used for sample acquisition for flatfield correction. z-stacks of 37 planes with a size of 200 nm were imaged, with no binning and dual-gain ¼ setting. For all images, z-stacks were imaged in the following order: first all laser lines were imaged before the next z-plane. The final pixel size was 0.108 µm x 0.108 µm. Exposure times (usually between 500 and 700 ms) and laser power varied between experiments due to different probe sets used in SABER-FISH. If not stated differently, images of the residual experiment were taken with no binning, dual gain ¼ settings and by using a silicon oil objective with correction collar (Plan Apo λS SR HP 100xC/1.45 Silicon DIC) in combination with silicon immersion oil 30cc (Nikon, #MXA22179). For collar correction of the silicon oil objective, fluorescent beads were prepared as described in (Isaac *et al*., 2024). Z-stacks with varying plain number to image full volumes of cells and sizes of 200 nm or 300 nm were taken. The final pixel size was 0.065 µm x 0.065 µm. RNase and DNase treated samples were imaged with a Plan Fluor 40x Oil DIC H/N2 objective in combination with immersion oil, 12-bit and no binning settings. The final pixel size was 332.8 µm x 332.8 µm.

### Image Analysis

#### Flatfield correction of images

For image analysis, the images were first background corrected in Fiji. Therefore, the average of 10 field of views of an empty well imaged with the same microscopic settings used to image the samples of each experiment were averaged. The sample image and the background image were corrected for the camera error by subtraction. Afterwards, the sample image was divided by the background. Next followed a z-projection of the z-stack using the sum of each slice in Fiji.

#### Segmentation of whole cells and RNAs

Segmentation of cells and RNA signal was done in CellProfiler. For whole cell segmentation, the whole range of intensities was chosen and the cells were segmented by hand based on the WGA staining. To segment the RNA signal, again the full intensity range was used. Next, the borders were enhanced using the Kirsch filter. To detect primary objects, a global threshold was used with the following settings: Minimum Cross-Entropy, threshold smoothing scale of 1.3488, threshold correction factor of 1, lower and upper bounds were set to 0 and 1, no log transformation was done before thresholding and intensities were used to distinguish between clumped objects and used to draw dividing lines. The size used for the smoothing filter for declumping as well as the minimum allowed distance between maxima was set to automatic calculation. Full resolution was used to determine local maxima. Holes in identified objects were filled after declumping.

The segmented RNA were assigned to the segmented cells and the area as well as the intensities for the segmented cells and the RNAs were extracted. Intensity extraction was performed based on the RNA channel. The data could then be analyzed in R. The extracted data of the segmentation, used for analysis as well as normalized data is shown in supplemental tables 2 and 3.

#### Calculating the cellular area occupied

To calculate the cellular area occupied, the area of each RNA object segmented of a cell was summed up and divided by the area of the parent cell. The distribution was plotted as a swarm plot to represent single cells and on top of those box plots were plotted to show the median. To test if there was a significant change an one-sided t-test was performed using the rstatix R package and the following settings: pairwise_t_test(NormArea ∼ Time, paired = FALSE, alternative = “less”, p.adjust.method = “none”), where NormArea is the fraction of cellular area occupied. Statistical results are summarized in supplemental table 4.

#### Single cell intensity analysis

For intensity analysis of single cells the integrated intensities measured of each RNA object in a single cell were summed and normalized to the cell area. Again, single cell data were blotted as swarm plots and bar plots.

### Image Manipulations

To represent images in figures, brightness and contrast was adjusted for the whole field of views in Fiji to represent the full spatial distribution of RNAs. This was necessary due to high differences in intensities between control and sample and in an image itself. Thus, images do not represent differences in intensities. Images were then cropped for single cells and zoom-ins using regions of interest. For figures showing WGA staining, a mean filter size 1 was used for the WGA channel. All other channels were only adjusted for brightness and contrast. Transmission electron microscopy images were just cropped to show regions of interest of different cells.

### Transmission electron microscopy

Cells were treated for 24 h with IMT1B, DMSO or for 5 h with EtBr. Cells were fixed in 2.5% glutaraldehyde, 1.25% paraformaldehyde and 0.03% picric acid in 0.1 M sodium cacodylate buffer pH 7.4 for 1 hour. The cells were then postfixed for 30 min in 1% osmium tetroxide (OsO4)/1.5% potassiumferrocyanide (KFeCN6), washed in water 3x and incubated in 1% aqueous uranyl acetate for 30 min followed by 2 washes in water and subsequent dehydration in grades of alcohol (5 min each; 50%, 70%, 95%, 2x 100%). Cells were removed from the dish in propyleneoxide, pelleted at 3000 rpm for 3min and infiltrated for 2 h to ON in a 1:1 mixture of propyleneoxide and TAAB Epon (TAAB Laboratories Equipment Ltd, https://taab.co.uk). The samples were subsequently embedded in TAAB Epon and polymerized at 60°C for 48 h.

Ultrathin sections (about 80 nm) were cut on a Reichert Ultracut-S microtome, picked up on to copper grids stained with lead citrate and examined in a JEOL 1200EX Transmission electron microscope or a TecnaiG² Spirit BioTWIN and images were recorded with an AMT 2k CCD camera.

### Transient transcription analysis

To measure transient transcription during recovery, cells were first treated for 5 hours with IMT1B as described above. The cells were washed three times with 1xPBS and grown in normal media for recovery. To label nascent RNAs, 4-thiouridine (4sU, Sigma T4509) was directly added to the cells to a final concentration of 500 µM at the last 10 min of each time point. Samples were harvested after 0, 1, 2 ,3 ,4 and 5 hours of recovery. Therefore, the media was removed and cells placed on ice. They were scraped directly into Quizol and the lysates were transferred into tubes. 2 mM DTT was added to each tube and the samples were heated for 3 min at 65°C and stored at -80°C for RNA extraction, followed by biotinylation and enrichment of labeled RNAs to be analyzed by MitoStrings.

### *4sU* labeling to distinguish between old and new RNA during recovery

To measure RNA abundance of old RNAs after stress release, cells were first treated with EtBr for 5 hours as described above. Afterwards, cells were washed three times with 1x PBS. Cells were then incubated in media supplemented with 500 µM 4sU for 2, 3, 4 or 5 hours. For time point 0 h, cells were harvested directly after EtBr treatment. Harvesting of cells was performed as described in the transient transcription section. The lysates could be used for RNA extraction and biotinylation. The labeled RNAs were bound onto streptavidin-beads and the flow-through fraction was used for analysis by MitoStrings.

### RNA extraction

RNAs of long-term treated cells and for RNA-seq were extracted as follows. After the respective treatment times, plates were put on ice. Cells were washed with ice cold 1xPBS and scraped into Quizol. Lysates were stored at -80°C until RNA extraction. Lysates were thawed at RT. 200 µl chloroform was added to 1 ml of sample, mixed by vortexing and incubated for 2 min at RT. Lysates were centrifuged for 15 min at 4°C and at 12,000 xg. The aqueous phase was transferred into a fresh tube without disturbing the interphase. RNAs were precipitated with 500 µl isopropanol and incubated either at RT for 10 min or at -80°C overnight. After a second spin at 4°C and 12,000 xg for 10 min the supernatant was discarded. The pellet was washed with 75% EtOH and pelleted again by centrifugation at 4°C and 7,500 xg for 5 min. The supernatant was discarded and the pellet dried for 5 to 10 min at RT. The RNAs were resuspended in 32 µl of RNAse-free water and the concentration was measured by Qubit and nanodrop. 40 ng of RNAs were used in MitoStrings measurements. For samples used to study the fate of old RNAs after stress release, 1 ng/µl unlabelled in vitro synthesized ERCC-00048 spike-in was added to the thawed cell lysates. In case of transient transcription analysis, 0.25 ng/µl of in vitro synthesized 4sU labeled ERCC-00136 spike-in (as described in (McShane *et al*., 2024)) was added additionally. RNAs were extracted as described and in these cases, RNA was resuspended in 50 µl of RNAse free water.

### RNA biotinylation and purification

A maximum of 40 µg of RNA per sample were used for the biotinylation reaction. Samples were brought to a volume of 70 µl and incubated at 60°C with 800 rpm rotation for 10 min. Afterwards samples were immediately incubated for not longer than 2 min on ice. RNAs were biotinylated in a reaction containing 1x labeling buffer (20 mM Hepes pH 7.4, 1 mM EDTA) and 0.5 mg/ml Biotin-MTS (Biotium, #90066) in 20% dimethylformamide (Sigma). The reaction mixture was incubated in a thermoblock at 800 rpm and 24°C for 30min in the dark. Next free biotin was removed. To this end, 2 ml phase-lock heavy gel tubes (5Prime, #2302830) were prepared by centrifugation at 14,000 xg for 1 min. The biotinylated RNA was diluted with water, transferred into the phase-lock tube and mixed with 1 volume of chloroform:isoamylalcohol (24:1) by manually shaking for 15 s. A centrifugation step for 5 min at RT and 16,000 xg followed. The upper phase was transferred into a fresh tube and the RNA was precipitated by addition of 0.1 volume of 5 M NaCl and 1 volume of isopropanol. The mixture was centrifuged for 40 min at 4°C and 20,000 xg. The supernatant was removed and the pellet washed with 75% EtOH, followed by a 5 min spin at 4°C and 7,500 xg. The supernatant was removed and the pellet was dried for 5 min at RT. The RNA was resuspended in 40 µl of RNAse-free water.

Samples could be stored at -80°C for enrichment. The biotinylated RNA was purified with streptavidin beads of the uMACS Streptavidin kit (Miltenyi Biotec #130-133-282). 3 µg RNA were mixed with streptavidin beads, incubated for 15 min at RT and subsequently loaded onto a uMACS column. The flow-through was loaded again onto the columns to increase biotin binding. The flow-through contained the unlabeled RNA fraction. The beads were washed three times with prewarmed (65°C) washing buffer (100 mM Tris pH 7.5, 10 mM EDTA, 1M NaCl and 0.1% Tween-20) and three times with washing buffer at RT. RNAs were eluted twice by incubation at 65°C in elution buffer (550 mM Tris, pH9, 10 mM EDTA, 150 mM TCEP). For transient transcription analysis the elution fraction, and to analyse old RNA, the flow-through fraction were purified using the miRNeasy Micro kit following the manufacturer’s instructions including the DNAse treatment (Qiagen, #217084). 30 ng RNA were used for MitoStrings analysis.

### MitoStrings

MitoStrings is a mitochondrial tailored NanoStrings measurement(Wolf and Mootha, 2014). According to the manufacturer’s protocol, RNAs (30 to 40 ng) were hybridized for 16 h at 67°C with the XT Tagset-24 (NanoString Technologies) and with DNA-probes specific for mitochondrial RNA (MitoStrings-probes were modified as in (Isaac *et al*., 2024)) in hybridization buffer (NanoString Technologies). Afterwards samples were loaded onto a nCounter Sprint Cartridge and quantified by use of the nCounter SPRINT Profiler (NanoString Technologies) of the Bauer core facility of Harvard or at the Manning lab (Harvard T.H. Chan School of Public Health). All raw counts are listed in supplemental table 5.

### Total RNA-sequencing

#### Sample collection

To analyze changes in OXPHOS expression, total RNA-seq was performed. HeLa cells were treated for 5 or 24 hours with EtBr or water as described before. Cells were harvested and the RNAs were extracted. The SMARTer Stranded Total RNA HI Mammalian kit (Takara #634873) was used to prepare sequencing libraries according to the manufacturer’s instructions. The libraries were measured on the NOVAseq (Illumina, San Diego, CA) by the Biopolymers Facility at Harvard Medical School.

#### Read Alignment

The provided fastq files were first concatenated to combine the data. Next, adapter sequences were trimmed as described in (McShane *et al*., 2024). The trimmed reads were filtered after mapping to rRNA using bowtie1(Langmead *et al*., 2009) with settings allowing for one internal and one 5′ mismatch. The residual reads were aligned to the GRCh38 human reference genome using STAR 2.7.3(Dobin *et al*., 2013) with the following parameters: --outSAMtype BAM SortedByCoordinate -- outReadsUnmapped Fastx --outFilterIntronMotifs RemoveNoncanonicalUnannotated -- outFilterMultimapNmax 100.

### Expression analysis

To analyze expression changes of OXPHOS genes, read counts were normalized to length and library size-normalized read counts. Read counts were summed over protein coding regions with the help of the R package featureCounts(Liao, Smyth and Shi, 2019) with the parameters: isGTFAnnotationFile = TRUE, useMetaFeatures = TRUE, countMultiMappingReads = FALSE, strandSpecific = 1, isPairedEnd = TRUE, nthreads = 4. GTF files used in this study are described here (Soto *et al*., 2022). The RPKM data and fold change can be found in supplemental table 6.

### Half-life calculations

To calculate half lifes, RNA abundance was measured in long-term transcription inhibition treatments by MitoStrings. RNA levels were measured after 0, 2, 5, 8 and 24 hours of transcription inhibition. The mean of three biological replicates (EtBr treated cells) or two biological replicates (IMT1B treated cells) of GAPDH normalized RNA counts were used for calculating half-lifes. 0, 2 and 5 hour measurements were used in linear regression with log transformation and time point 0 h was set as 100%. The slope was used to calculate the half lifes: (log10(0.5)-log10(1)) / slope. The same procedure was performed for late half-lifes. Here, the 5 hours measurements were set as 100%. For NDUFA7, we used for early half-lifes only 0h and 5h as the 2 hour time point increased and led to negative half-lifes.

### Use of AI

chatGTP was used to help formulate R code for half-life analysis, to extract statistics and to write functions for image analysis. Claude was used to edit and improve the text.

